# Assessment of pre-clinical liver models based on their ability to predict the liver-tropism of AAV vectors

**DOI:** 10.1101/2022.09.28.510021

**Authors:** Adrian Westhaus, Marti Cabanes-Creus, Kimberley L. Dilworth, Erhua Zhu, David Salas Gómez, Renina G. Navarro, Anais K. Amaya, Suzanne Scott, Magdalena Kwiatek, Alexandra L. McCorkindale, Tara E. Hayman, Silke Frahm, Dany P. Perocheau, Bang Manh Tran, Elizabeth Vincan, Sharon L. Wong, Shafagh A. Waters, Laurence O. W. Wilson, Julien Baruteau, Sebastian Diecke, Gloria González-Aseguinolaza, Giorgia Santilli, Adrian J. Thrasher, Ian E. Alexander, Leszek Lisowski

**Affiliations:** Translational Vectorology Research Unit, Children’s Medical Research Institute, Faculty of Medicine and Health, The University of Sydney, Westmead, Australia; Great Ormond Street Institute of Child Health, University College London, London, UK; Gene Therapy Research Unit, Children’s Medical Research Institute and Sydney Children’s Hospitals Network, Faculty of Medicine and Health, The University of Sydney, Westmead, Australia; Universidad de Navarra, CIMA, Gene Therapy and Regulation of Gene Expression Department, Avenida Pío XII, Pamplona, Spain. IdiSNA, Instituto de Investigación Sanitaria de Navarra; Australian e-Health Research Centre, Commonwealth Scientific and Industrial Research Organisation, New South Wales, Sydney, Australia; Military Institute of Hygiene and Epidemiology, The Biological Threats Identification and Countermeasure Centre, Puławy, Poland; Inventia Life Science Pty Ltd, Sydney, Australia; Stem Cell Technology Platform, Max Delbrück Centrum for Molecular Medicine, Berlin, Germany; Genetics and Genomic Medicine Department, Great Ormond Street Institute of Child Health, University College London, UK; Metabolic Medicine Department, Great Ormond Street Hospital for Children NHS Foundation Trust, London, UK; National Institute of Health Research Great Ormond Street Hospital Biomedical Research Centre, London, UK; Molecular Oncology Group and Victorian Infectious Diseases Reference Laboratory, The University of Melbourne, The Peter Doherty Institute, Melbourne, Australia; Molecular and Integrative Cystic Fibrosis (miCF) Research Centre, University of New South Wales and Sydney Children’s Hospital, Sydney, Australia; School of Biomedical Sciences, Faculty of Medicine, University of New South Wales, Sydney, Australia; Department of Respiratory Medicine, Sydney Children’s Hospital, Faculty of Medicine, University of New South Wales, Sydney, Australia; Applied BioSciences, Faculty of Science and Engineering, Macquarie University, New South Wales, Sydney, Australia; Discipline of Child and Adolescent Health, Faculty of Medicine and Health, The University of Sydney, Sydney, New South Wales, Australia; Vector and Genome Engineering Facility, Children’s Medical Research Institute, Faculty of Medicine and Health, The University of Sydney, Westmead, Australia; Laboratory of Molecular Oncology and Innovative Therapies, Military Institute of Medicine, Warszawa, Poland

## Abstract

The liver is a prime target for *in vivo* gene therapies using recombinant adeno-associated viral vectors (rAAV). Multiple clinical trials have been undertaken for this target in the past 15 years, however we are still to see market approval of the first liver-targeted AAV-based gene therapy. Inefficient expression of the therapeutic transgene, vector-induced liver toxicity and capsid, and/or transgene-mediated immune responses reported at high vector doses are the main challenges to date. One of the contributing factors to the insufficient clinical outcomes, despite highly encouraging preclinical data, is the lack of robust, biologically- and clinically-predictive preclinical models. To this end, this study reports findings of a functional evaluation of six AAV vectors in twelve preclinical models of the human liver, with the aim to uncover which model is the most relevant for the selection of AAV capsid variant for safe and efficient transgene delivery to primary human hepatocytes. The results, generated by studies in models ranging from immortalized cells, iPSC-derived and primary hepatocytes, and primary human hepatic organoids to *in vivo* models, increased our understanding of the strengths and weaknesses of each system. This should allow the development of novel gene therapies targeting the human liver.

## Introduction

Recombinant adeno-associated viral (rAAV) vectors are versatile delivery tools composed of a single-stranded DNA genome flanked by 145-bp inverted terminal repeats (ITRs), packaged within an icosahedral protein capsid. The adeno-associated virus (AAV), from which this vector system was derived, is a non-pathogenic helper-dependent parvovirus with multiple naturally occurring serotypes, including the prototypical serotype 2 (AAV2).^1,2^ AAV2 was the first variant to be vectorized and is the best understood serotype, still used in many studies to date.^3,4^ The structure and amino acid sequence of the non-enveloped AAV capsid is the main determinant of tropism.^5^ Therefore modifying the viral capsid has been used as a strategy to target specific cell types and organs for therapeutic applications.^6^

Clinical success of gene therapy trials using AAV vectors has led to the authorization of products for three indications to date: RPE65-associated retinal dystrophy (AAV2 capsid, Luxturna™),^7^ spinal muscular atrophy (SMA) (AAV9 capsid, Zolgensma™),^8^ and lipoprotein lipase deficiency (AAV1 capsid, Glybera™; no longer available).^9^ While these AAV-based products target different organs (the eye, the CNS, or the muscles, respectively), therapies targeting disorders of other organs, such as the liver, have not reached market approval to date.

The liver is an important clinical target for gene therapies because of its key role in metabolism and homeostasis. Most of the experience in liver gene transfer using AAV vectors has been obtained in clinical trials for two coagulation disorders: Hemophilia A and B.^10,11^ Data from these trials have been encouraging and showed a relatively good safety profile. However, clinical studies conducted to date pointed out several challenges that need to be overcome to facilitate approval of therapeutic products for these diseases and expanding AAVs as therapeutics for other liver disorders. These challenges include the activation of the immune system following vector administration, unexpectedly low efficiency of targeting human hepatocytes *in vivo* and liver toxicity associated with the administration of high vector doses.^12^

The development of immune responses towards the therapeutic vector, resulting in the reduction of transgene expression, was first observed in a Phase I/II study that utilized AAV2 to express Factor IX to treat Hemophilia B.^13^ AAV2 is endemic to the human population and thus many patients who could benefit from AAV2-based therapeutics have developed immunity to this serotype during their lifetimes, including six out of seven patients enrolled in the first systemic clinical AAV2 study. Elevation of liver transaminases and CD8^+^ T-cells against AAV were detected following vector administration through the hepatic artery.^13,14^ Increase in Factor IX levels was only detected in two patients and it was transient, a striking contrast to data obtained in preclinical studies in mice and non-human primates (NHPs), which showed long-term transgene expression.^15^ While overall disappointing, the study confirmed the relative safety of rAAV-mediated liver gene transfer.^16^ Critically, however, the activation of the immune system was not observed in any of the studies performed in animal models, highlighting the limitations of the model systems used to develop and validate new AAV-therapeutics prior to clinical implementation.

Subsequent liver-targeted trials utilized other serotypes, such as AAV8^17^ and AAV5,^18^ both selected based on preclinical data in mice and NHPs.^19,20^ In a pivotal trial sponsored by St. Jude Children’s Research Hospital, a self-complementary AAV8 vector encoding Factor IX was administered to seronegative patients at three different doses.^21^ Although long-term clinical efficacy was achieved, an early increase in liver transaminases was observed in the high dose cohort.^22^ Immune adverse events in liver clinical trials can generally be controlled by the administration of corticosteroids to prevent elimination of transduced cells and thus therapeutic transgene expression. Nevertheless, prevalence of neutralizing antibodies reduces the pool of patients that may be able to benefit from novel experimental therapies. However, clinical studies suggest that anti-AAV5 antibodies do not preclude successful liver transduction with AAV5-based vectors.^23^ This vector serotype has also been used to deliver Factor IX at relatively high vector doses (2×10^13^ vg/kg) with no significant T-cell mediated inflammation,^24,25^ although it must be considered that this serotype is also less efficient than others at transducing hepatocytes in animal models.^19^

Despite these current limitations, it remains clear that vector serotype selection plays a crucial role in obtaining the desired clinical outcomes. AAV serotypes used in clinical studies are selected based on data generated in preclinical, frequently murine, models. However, clinical data obtained with AAV2, AAV5 and AAV8 clearly indicate that AAV liver-tropism, which can be species-specific,^20^ can differ significantly between the murine or NHP preclinical models and human patients.^11^

One way to overcome this issue has been using “humanized” models to functionally evaluate the existing AAV variants for their ability to transduce human hepatocytes *in vivo*. The same models can also be used to identify novel capsids with improved liver-tropism. To this end, bioengineered capsids with high tropism towards human hepatocytes, such as AAV-LK03^26^ and AAV-NP59^27^ were developed in xenografted *Fah^-/-^/Rag2^-/-^/Il2rg^-/-^* (FRG)^28^ mice demonstrating the validity of this approach. The shuffled capsid AAV-LK03, bearing high sequence similarity to the natural serotype AAV3b,^1,29^ has been the first non-natural AAV to be used in clinical studies and led to stable therapeutic levels of Factor VIII expression in 16 out of 18 Hemophilia A patients.^30^ Two patients treated at the highest dose in this study lost FVIII expression likely as a result of immune activation against the vector.^30^ AAV-NP59, a highly functional variant selected in humanized FRG, which differs from prototypical AAV2 at eleven amino acids only,^31^ is yet to be tested in the clinic.

The clinical progress is further affected by our inability to directly compare clinical data obtained from liver gene transfer studies using different AAV vectors not only due to the differences imposed by the individual AAV variants used, but also due to the lack of consistency regarding preclinical models used. Furthermore, vector manufacturing protocols, quantification methods, and transgene expression cassettes differ between the individual studies conducted to date.^32^ Finally, patient-to-patient variability adds yet another level of complexity and “noise” in clinical data, making it very difficult to draw conclusions on how individual vector performance compares to one another.

The choice of the therapeutic cassette is dictated by the specific indication being targeted, and the patient-to-patient differences are impossible to overcome. However, the lack of consistency in the preclinical models used to select the rAAV variant needs to be addressed to determine critical decisions such as selection of the vector type and clinical vector dose. Advances in this respect will enable the successful development of novel effective and safe gene therapeutics.

With this in mind, we set out to compare AAV-based gene transfer efficiency targeting the liver in twelve frequently used preclinical models, including *in vitro* models, such as hepatic cell lines, human induced pluripotent stem cell (hiPSC)-derived hepatocytes (iHeps), and adult stem cell-derived hepatic organoids. However, the focus was on *ex vivo* models, such as 2D and 3D primary NHP and human hepatocytes cultures, and *in vivo* models, including murine and human hepatocytes in xenograft mice and NHPs. To ensure the data obtained was not unique to the specific AAV variant used in the study, we performed the studies using six natural and bioengineered AAV variants, including four that have previously been utilized in liver-directed clinical studies. Moreover, to increase quality and impact of the study by minimizing experimental noise, we used a well-characterized barcode approach, which allowed us to compare the individual vectors at the cell entry and transgene expression levels in parallel using next-generation sequencing (NGS) in each of the models.^33–35^ This study was aimed to understand the differences between the models and what will be required to address future preclinical translation. On top of the expected differences between the models in respect to their serological and immunological properties, we identified surprising differences between NHP and human hepatocytes that might explain some of lower-than-expected outcomes of clinical trials to date.

## Material and Methods

### Cell culture conditions and cell origins

AAV production and anti-AAV neutralization assays were performed in a human embryonic kidney (HEK) 293T cell line (ATCC, Cat# CRL-3216). HEK293T cells were cultured in Dulbecco’s Modified Eagle’s Medium (DMEM, Gibco, Cat# 11965) supplemented with 10 % fetal bovine serum (FBS, Sigma-Aldrich, Cat# F9423), 1× Pen Strep (Gibco, Cat# 15070), and 25 mM HEPES (Gibco, Cat# 15630). Human hepatocellular-carcinoma 7 (HuH-7) and hepatocellular carcinoma HepG2 cells were provided by Dr Jerome Laurence (The University of Sydney) and Prof. Ian E. Alexander, respectively, and cultured in DMEM supplemented with 10 % FBS, 1× Pen Step and 1× Non-essential amino acids (Gibco, Cat# 11140).

### Cell transductions

Transductions were performed as previously published. ^35,36^ Briefly, AAVs were added the indicated amount of rAAV capsid or vector mix to the cells, changing media after six-to-eight hours and harvesting cells for DNA/RNA or flow cytometry processing 72 hours after rAAV exposure, unless otherwise specified.

### Origin and culture of iHEP

Frozen hiPSC-derived hepatocytes (from ChiPSC18) were purchased at Takara Bio Europe AB, thawed, plated, and maintained according to the manufacturer’s instructions in media included in the kit (Cellartis Enhanced hiPS-HEP v2; Cat# Y10134). Briefly, 8.2 × 10^5^ cells/well were thawed and seeded in a 24-well plate, exposed to AAVs after five days in culture and harvested seven days after exposure.

### Primary human hepatocytes in 2D- and 3D-culture

Human (Lonza, HUM4198) and rhesus macaque (Lonza, Cat# MKR103) primary hepatocytes were used for 2D and 3D culture systems. For 2D culture, 24-well plates were coated with collagen for 45 minutes. After washing the coated wells with PBS, primary hepatocytes were seeded 400,000 cells per well in complete hepatocyte plating media (HCM kit, Lonza, Cat# CC-3198). After four hours of incubation at 37 °C and 5 % CO_2_, media was removed and gently covered with hepatocyte maintaining media (HCM kit, Lonza, Cat# CC-3198) containing 0.25 mg/mL Matrigel (Corning, Cat# 354234) and incubated at 37 °C for 90 minutes. The AAV mix was added diluted in hepatocyte basal media (HBM, Lonza, Cat# CC-3199). Media was changed daily for three days, until harvest using Cell Recovery Solution (BD, Cat# 354253) and processing for NGS.

The 3D printed hydrogels containing human and rhesus hepatocytes were generated using a Rastrum Cell Printer (Inventia, Sydney, Australia). The cell printing followed the manufacturer’s instructions and previously published protocols.^37,38^ Briefly, hepatocytes were thawed, resuspended in crosslinker solution (Inventia) and printed in 96-well plates at 8,000 cells per well. Cells were maintained in HBM and AAVs were added as described above. Cells were harvested as whole printed gels and processed for NGS.

### Primary human and simian hepatocytes engrafted into FRG mice

Cynomolgus and Rhesus macaque primary hepatocytes were purchased from Lonza (Cat# MKC118 and Cat# MKR103). Human hepatocytes were also purchased from Lonza (Cat# HUM181971) and corresponded to a 15-month-old donor.

The engraftment procedure was performed as previously published.^28,39^ Mouse studies were supported by the BioResources Core Facility at Children’s Medical Research Institute. All animal care and experimental procedures were approved by the joint Children’s Medical Research Institute and The Children’s Hospital at Westmead Animal Care and Ethics Committee. The FRG mice were housed, treated, and sacrificed following previously published methods.^39^

### Liver organoid transduction

Human liver organoids were generated as previously described^40^ and used for research purposes with approval from the human ethics committee of School of Biosciences, University of Melbourne (ethics number 1851272). To carry out transduction with AAV, the Matrigel-supported planar infection method was adapted.^41^ Briefly, mature liver organoids embedded in Matrigel domes were isolated by dispersing Matrigel with cold basal media. After centrifugation, the medium was discarded, and the organoid pellet was suspended with expansion medium containing 10 μM Y-27632 (Selleckchem, Cat# S1049). The AAV cocktail was added to the medium at the described dose, and then the organoid–AAV mixture was transferred to a 24 well plate pre-coated with 80 μl of 75 % Matrigel. The organoid-AAV mixtures were incubated at 37 °C with 5 % CO_2_ for 12 hours in a planar manner after which organoids on the Matrigel surface were collected, transferred to a tube, centrifuged, and supernatant discarded. The organoids were then washed with cold basal media followed by centrifugation. After the final wash and centrifugation, the organoids were returned to 3D culture by resuspending in Matrigel and seeding at 50 μl per well into 24-well plates. Once the Matrigel had set, 450 μl of expansion culture medium (Supplementary Table 1) was added and then replaced every other day for seven days. To harvest, the organoids were suspended in cold basal media after the medium was discarded, pelleted by centrifugation, and then snap frozen until processing for DNA and RNA extraction. All centrifugations were carried out at 400 × g at 4 °C for five minutes.

### PCR reactions

Standard and Illumina amplicon-seq NGS polymerase chain reactions (PCRs) were performed using Q5 [NEB, Cat# M0491], dNTPs [NEB, Cat# N0447] and primers (all Sigma-Aldrich, Supplementary Table 2) and were performed strictly following previously published protocols.^35^

### AAV production

All AAV capsids used in this study were produced using polyethylenimine (PEI) transfection of the LSP-GFP-barcode (LSP-BC) transgene cassette,^31,36,42,43^ as well as adenovirus and Rep2-Cap2/3/5/8/LK03/NP59 helper plasmids using previously described methods.^36^ The resulting cell lysates and purified media were either purified using iodixanol gradients (all experiments apart from NHP) or CsCl ultracentrifugation (for NHP vectors) following the previously published protocols, respectively.^26,35^ AAV titers were established using quantitative PCR and GFP primers following previously published methods.^35,44^ The capsids were then mixed at an equimolar ratio as previously described.^35^

### Neutralization Assays

For experiments using the NHP, neutralization assays determining the anti-capsid antibodies for the different AAVs for all indicated timepoints were performed after the following protocol. At day 1, HEK293T cells were seeded into a 96-well plate at a density of 10^4^ cells/well. At day 2, NHP sera were diluted in DMEM supplemented with 2 % FBS in a total volume of 100 μL, beginning with a 1:5 dilution followed by dilution series of 1:3 and mixed with a dose of 10^4^ vector genomes per cell (vg/cell) of the corresponding AAV serotype coding for luciferase that were incubated for 2 h at 37 °C. The mix was subsequently used to transduce target cells. Each serum dilution was tested in duplicate. Negative controls of non-transduced cells, as well as positive controls of cells transduced without AAVs not preincubated with NHP serum were included in each plate. The transduced cells were incubated for 48 h before quantification of luciferase activity was performed. Light emission from each well was measured in photons/sec. The NAb titer was calculated using the highest dilution for which the percentage of light emission was 50% of positive control without serum.

For antibody determination in human serum samples, the assay was performed as previously described.^45^ Briefly, HuH-7 cells were incubated with heat-inactivated sera at dilution starting from 1:5, continuing in 2-fold serial dilutions to 1:1280. The diluted serum samples were incubated for one hour at 37 °C with the individual AAV capsids diluted in an equal volume of DMEM. rAAV vectors were incubated at the same concentration to reach a predetermined final variant-dependent dose into a 100 μL final volume for transduction. The appropriate dose used was 6,000 vg/cell for AAV2, 2,000 vg/cell for AAV3, 40,000 vg/cell for AAV5, 25,000 vg/cell for AAV8, 2,000 vg/cell for AAV-LK03 and 4,000 vg/cell for AAV-NP59. Pooled human serum samples were used as a positive control. Quantification of GFP positive cells was performed by mean fluorescence using flow cytometry 72 hours after rAAV exposure. A 1:5 dilution of serum reducing the vector transduction by 50% or more was considered positive. The highest positive serum dilution determined the neutralizing antibody titer.

### Human serum samples

Human serum samples from the Immunology laboratory, Great Ormond Street Hospital for Children NHS Foundation Trust, London, UK were anonymously analyzed following the guidelines of the Royal College of Pathologists.

### Non-human primate work

Animal procedures were approved by the ethical committee for animal testing of the University of Navarra and by the Department of Health of the government of Navarra (Comité de Etica para la Experimentación Animal code: 038/15) and performed according to the guidelines from the institutional ethics commission. Animal welfare checks were performed by animal care staff twice daily.

A young adult male *Macaca fascicularis* NHP was tested negative for anti-AAV NAbs for AAV2, AAV5 and AAV-LK03 and with low seropositivity against AAV3, AAV8 and AAV-NP59.

On day 0, the NHP was anesthetized, blood was drawn to obtain serum and the animal was subjected to the immunoadsorption process (described in Salas *et al.*,^46^). Within the following 30 min after immunoadsorption, the vector was infused via the saphenous vein over ten minutes. Blood was collected at one hour, 24 hours and seven days after administration of the vector. At day 7, the animal was euthanized, and different organs and tissues were collected for further analysis.

### DNA and RNA extraction from cells and tissue samples

DNA and RNA were isolated from the cell pellets from the *in vitro* and *ex vivo* experiments using the AllPrep DNA/RNA Mini Kit (QIAGEN, Cat# 80204) following the manufacturer’s instructions.

DNA from the mouse, NHP and human hepatocytes from xenograft FRG mice and NHP tissues were isolated using phenol-chloroform extraction after proteinase K digestion following previously published protocols.^35^ RNA from the mouse, NHP and human hepatocytes from xenograft FRG mice and NHP tissues was extracted using the TriReagent (Sigma, Cat# T9424)-chloroform protocol previously published.^35,36^

### Reverse transcription of extracted RNA

Clean-up of extracted RNA was performed using TURBO DNase (Invitrogen, Cat# AM1907) twice for one hour, followed by incubation with DNAse inactivation reagent following the manufacturer’s instructions. The DNase-treated RNA was then used for cDNA synthesis using the SuperScript IV First-Strand Synthesis System (Invitrogen, Cat# 18091050) following the manufacturer’s instructions using 2 μM of WPRE-binding primer (WPRE_R, Supplementary Table 2) to specifically synthesize the barcoded ssAAV-LSP-GFP-BC-WPRE cDNA.

### Next-Generation Sequencing (NGS)

The NGS amplicons were prepared and analyzed as previously published.^35,36^

### Normalization of NGS reads

NGS data obtained from all samples were normalized to the barcode contribution of the respective input (as indicated above). Read counts for each sample and each variant were multiplied by the variant specific ‘normalization co-efficient’ of the respective input, which was calculated as follows:

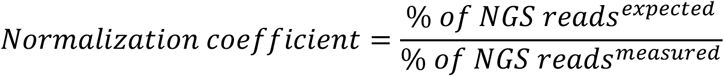

## Results and Discussion

### Study design

To compare the performance of different AAV variants in preclinical models of the human liver, we created an equimolar mix of the following six AAV variants: AAV2, AAV3b, AAV5, AAV8, AAV-LK03 and AAV-NP59; encoding barcoded expression cassettes compatible with NGS-based readout at the cell entry (DNA) and transgene expression (RNA/cDNA) levels.^31,35,42^ AAV2, AAV5, AAV8 and the bioengineered AAV-LK03 were chosen as clinical data from liver-targeted human studies was publicly available.^11^ AAV-NP59 and AAV3b were included because they were the bioengineered variant of AAV2 and the closest natural relative of AAV-LK03, respectively (Supplementary Fig. 1).

The mix of the six AAV candidates was used to transduce all different preclinical models evaluated in this study, while the individual AAV candidates underwent seroprevalence studies using individual human sera. The functional data obtained was compared to publicly available data on AAV performance in clinical trials as well as between the models to identify differences (Supplementary Fig. 1).

### *In vitro* and *ex vivo* models of the human liver

The simplest models of human hepatocytes are based on cell lines derived from hepatocellular carcinoma, such as HuH-7, or hepatocellular blastoma, such as HepG2 cells.^47,48^ These lines are immortalized and self-renewing, allowing for low-cost high-throughput experimentation using standard laboratory equipment and reagents. To evaluate both cell types for their potential to serve as biologically predictive models of human primary hepatocyte transduction, cells were transduced at a dose of 1,000 and 200 vg/cell and were harvested 72 hours after AAV exposure.

Cell pellets were processed for DNA and RNA/cDNA, which allowed us to evaluate relative vector performance at the cell entry and transgene expression levels, respectively (Supplementary Fig 2). The NGS results showed that AAV2 outperformed the other variants at the DNA and RNA levels in both HuH-7 and HepG2 cells. Interestingly, while AAV-LK03 and AAV3b performed similarly at the cell entry level (DNA) in HuH-7 cells, AAV-LK03 outperformed AAV3b at the transgene expression level (RNA). In HepG2 cells, AAV-LK03 and AAV3b performed similarly but were less efficient at both DNA and RNA/cDNA levels than AAV5. The strong performance for AAV2 in those cell lines was expected based on data from HuH-7 cells we reported previously.^35^

Next, we studied the six vectors in hiPSC-derived hepatocytes (iHeps) and adult stem cell-derived ductal organoids. It quickly became apparent that the transcriptional dominance of AAV2, as shown in the immortalized cell lines, did not fully translate to iHeps. AAV2 was still able to enter the cells at the highest efficiency at the DNA level, followed by AAV-LK03, AAV3b, AAV5, AAV-NP59 and AAV8 (Fig. 1b), but at the transgene expression level, AAV-LK03 outperformed all other vectors (Fig. 1c).

**Figure 1.**
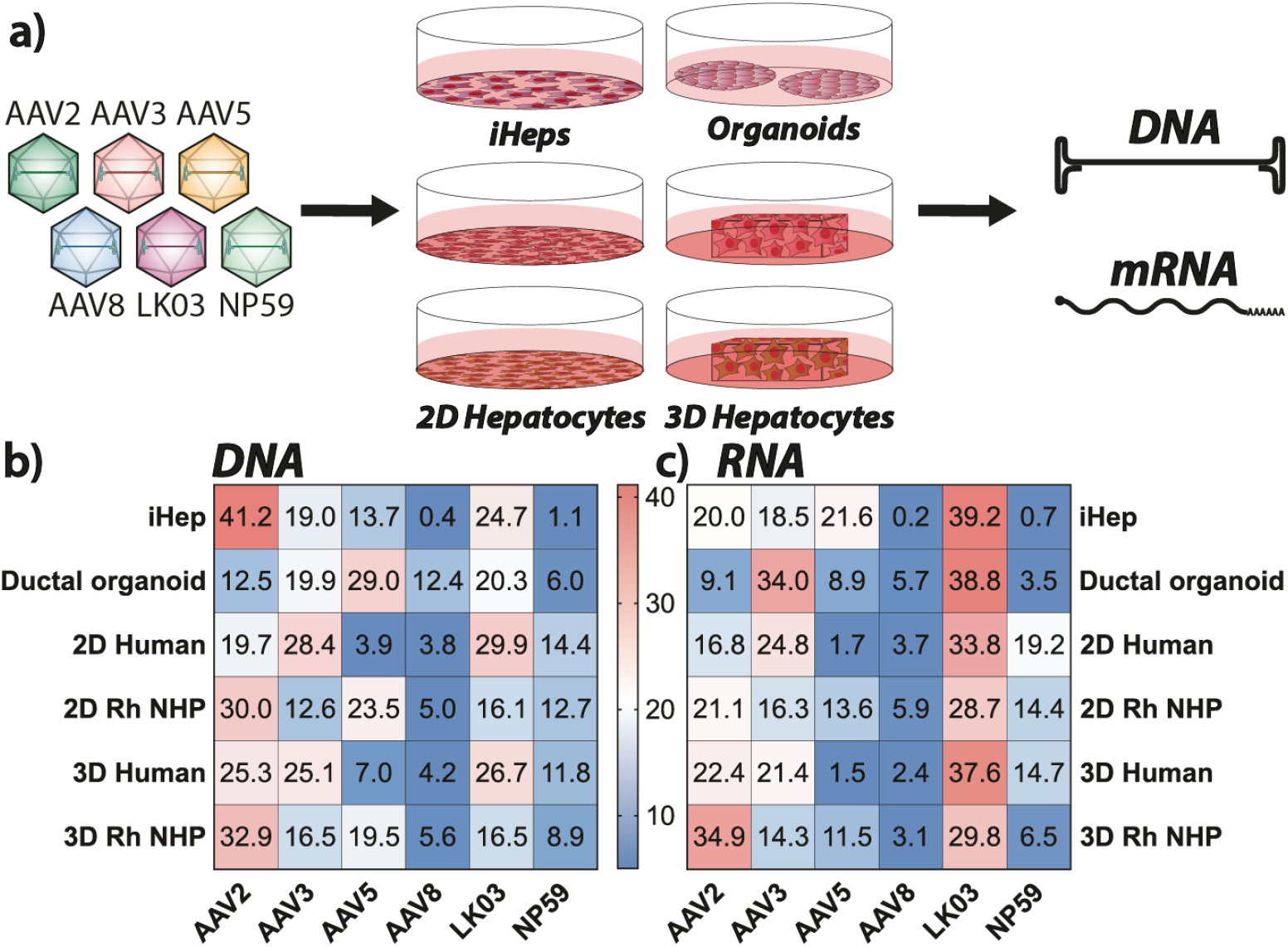
*Ex vivo* results in human and NHP liver models. (**a**) Schematic of transduction of indicated *in vitro* and *ex vivo* models. (**b**) NGS read contribution (%) for each AAV from extracted DNA. (**c**) NGS read contribution (%) for each AAV from mRNA-derived complementary DNA.

Using adult stem cell-derived ductal organoids consisting of liver ductal progenitor cells,^49^ we found AAV5 to be most effective at entering cells (29 % of NGS reads at the DNA level) (Fig. 1b). However, the efficient entry of AAV5 did not lead to efficient transgene expression, where only 8.9 % of the NGS reads from RNA/cDNA could be attributed to this variant. Instead, AAV-LK03 (38.8 %) and AAV3b (34 %) were most efficient at transgene expression in the organoids (Fig. 1c), despite less efficient entry (20.3 % and 19.9 % DNA NSG reads for AAV-LK03 and AAV3b, respectively). The observed decrease in the performance of AAV2 might indicate reduced importance of binding to heparan sulphate proteoglycan (HSPG) as a critical step for cell entry.^42^

To understand how the previous findings translate to primary cells *ex vivo,* human and rhesus monkey *(Macaca mulatta)* hepatocytes were seeded in 2D culture as well as 3D-printed in a hydrogel.^50^ The 2D cultured hepatocytes from human and NHP were exposed to three different AAV doses (1000, 500, and 200 vg/cell) and harvested three days after exposure to AAV. The hepatocytes of human origin were most efficiently entered by AAV-LK03 and AAV3b, followed by AAV2, AAV-NP59, AAV5 and AAV8 (Fig. 1b). In contrast, rhesus monkey primary hepatocytes were most efficiently entered by AAV2, followed by AAV5, AAV-LK03, AAV-NP59, AAV3b and AAV8 (Fig. 1b), showing a marked difference compared to transduction observed in the human hepatocytes. However, at the level of transgene expression, AAV-LK03 was the most efficient variant in both species. In human hepatocytes, it was followed by AAV3b, AAV-NP59, AAV2, AAV8 and AAV5, while in simian hepatocytes it was followed by AAV2, AAV3b, AAV-NP59, AAV5 and AAV8 (Fig. 1c).

As the next step, we wanted to transduce the same cells, but 3D-printed in a hydrogel substrate.^37^ However, before doing so, we evaluate the potential effect of the 3D culturing system on primary hepatocytes by performing RNAseq using cells before and after 3D culture (Supplementary Fig 3). The results showed that while expression of most genes did not change substantially, several genes underwent upregulation and downregulation after 3 days of the 3D culture (Supplementary Fig. 3b, Supplementary Table 3). For the human hepatocytes, the most upregulated gene was a long non-coding RNA (PCAT1) with the proposed function of regulating genes implicated in increased cell proliferation, migration, and invasion.^51,52^ Due to a lack of comprehensive annotation coverage of the rhesus monkey genome, the RNAseq data recovered for this species was less informative than the human data.

Hepatocytes in 3D hydrogel cultures were exposed to a dose of 500 vg/cell and cells were harvested and processed for analysis three days later. Results obtained showed number of differences compared to results obtained with conventional 2D cultures. Consistent with the 2D cultures, AAV-LK03 entered 3D-printed human hepatocytes with the highest efficiency among the vectors tested but was more closely followed by AAV2 and AAV3b. AAV-NP59, AAV5 and AAV8 performing substantially less efficiently. In the 3D-printed simian hepatocytes, AAV2 was found to be the most effective variant at cell entry, followed by AAV5, AAV3b and AAV-LK03, with AAV-NP59 and AAV8 being the weakest performers (Fig. 1b). At the transgene expression level (RNA) AAV2 gained in function in both human and rhesus cells when comparing to 2D cultures.

The data from 2D vs. 3D culture systems as well as human vs. NHP hepatocytes led to interesting observations. While AAV2 worked in 2D and 3D cultures of both species, there was a substantial increase in AAV2 performance in the 3D culture system over the 2D culture. This observation could be explained with strong reliance on HSPGs in the 3D printed hepatocytes, a question we explored further in subsequent studies. Furthermore, the data showed that AAV5 had a markedly higher performance in NHP cultured hepatocytes than in human hepatocytes, which might explain why clinical trials using AAV5 need very high vector doses and have a lower efficacy than NHP data would suggest.^24,25^

Another upregulated gene that is potentially relevant for AAV biology was glypican proteoglycan 6 (GPC6), a cell surface protein known to harbor HSPGs.^53^ This could explain the increased transduction by AAV2 and decreased transduction by AAV-NP59, which has lower affinity to HSPG (Fig. 1),^31^ in the 3D printed human hepatocytes compared to the conventionally cultured human hepatocytes.

In summary, in the simplest models of human hepatocytes, the immortalized cancer cell lines, AAV2 performed substantially better than all other variants tested. This performance decreased in stem cell-derived models and primary cells, where the performance of the bioengineered variant AAV-LK03 improved in both human and simian cells. The bioengineered AAV-NP59 also seemed to have improved in primary cells of human origin when compared to its performance in immortalized cells and stem cell-derived models. One of the most interesting observations was the relatively variable performance of AAV5 across the models tested. The performance was low in HuH-7 cells and primary human hepatocytes, while a relatively high performance was observed in HepG2 cells, stem cell derived iHeps and ductal organoids as well as primary NHP hepatocytes. This might indicate that AAV5 (the most distantly related AAV capsid of the ones chosen for this study) might utilize distinct cell entry and transduction mechanism.

### Xenograft *in vivo* models of human and non-human primate livers

Having studied the six vectors in several *in vitro* and *ex vivo* models, we next wanted to evaluate one of the commonly used *in vivo* xenograft model of the human liver, namely the FRG mouse.^28^ To facilitate the comparison to the data obtained from the *ex vivo* studies, we used primary human hepatocytes and primary rhesus hepatocytes, as in the studies in the 2D and 3D cultures, to engraft livers of female FRG mice. Additionally, we included primary hepatocytes from the cynomolgus monkey *(Macaca fascicularis).*

Thus, we generated humanized FRG (hFRG) and two types of “monkeynized” FRG, RhFRG and CyFRG, based on rhesus and cynomolgus origin of cells, respectively. Animals were allowed to repopulate to a replacement index ranging from 15 to 70 % and were subsequently systemically injected with the equimolar mix of the six barcoded AAV vectors (Fig. 2a). Livers were harvested seven days after transduction. Analysis of vector function at the DNA (cell entry) level was performed in sorted GFP^+^ and unsorted (total “bulk” fraction) human and simian cells, whereas analysis at the RNA level (transgene expression) was only performed on human and simian cells sorted based on the vector encoded GFP marker (GFP^+^).

**Figure 2.**
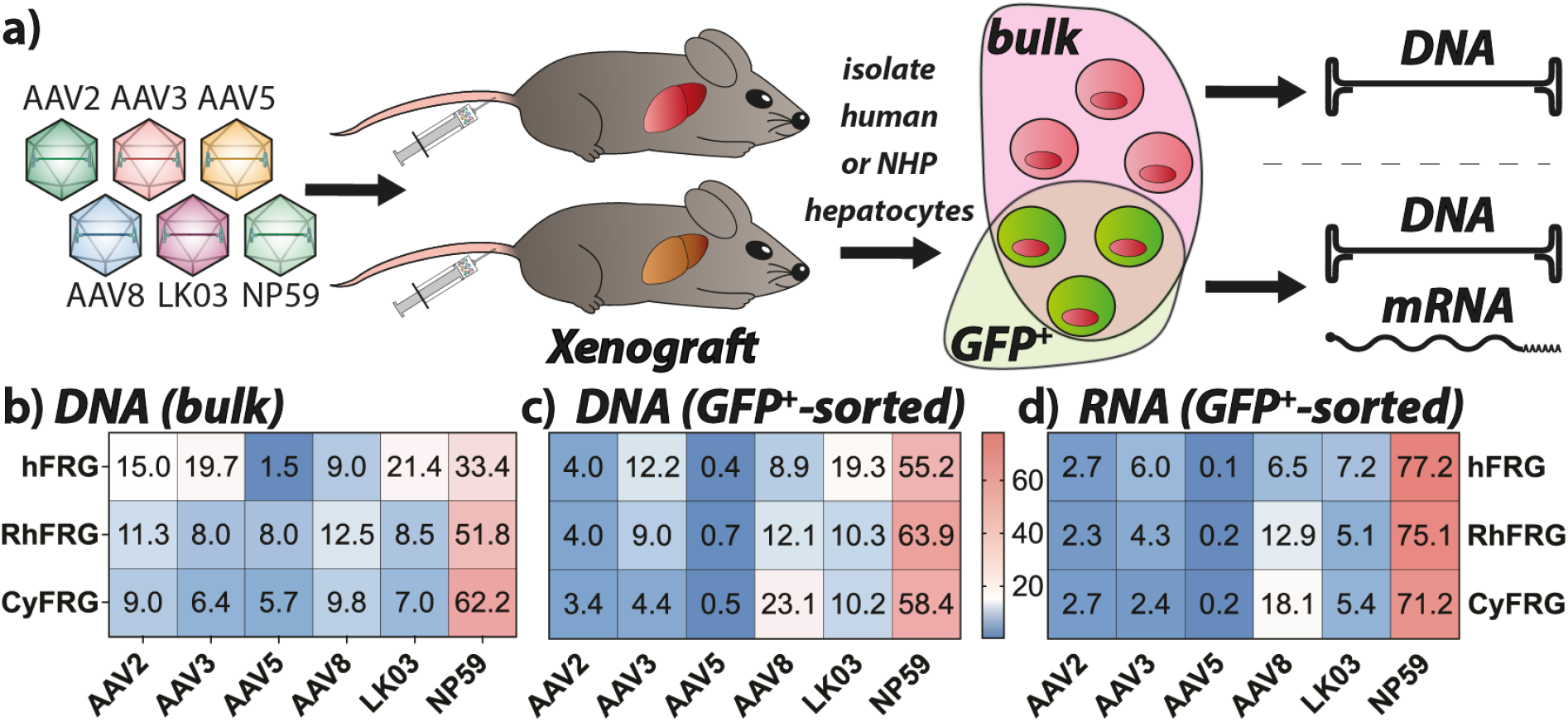
NHP and human xenograft *in vivo* results. (**a**) Schematic of transduction of engrafted FRG mice. (**b**) NGS read contribution (%) for each AAV from extracted xenograft bulk DNA. Cells were sorted for xenograft species only. (**c**) NGS read contribution (%) for each AAV from extracted eGFP-positive xenograft hepatocyte DNA. Cells were sorted for xenograft species as well as eGFP expression where indicated. (**d**) NGS read contribution (%) for each AAV from xenograft mRNA-derived complementary DNA. Cells were sorted for xenograft species as well as eGFP expression.

At the cell entry level, AAV-NP59 was the most effective variant in all three xenograft models irrespectively of analysis being performed on GFP^+^-sorted or bulk cells (Fig 2b-c). In bulk human cells, AAV-NP59 was followed by AAV-LK03 / AAV3b, AAV2, and AAV8, with AAV5 being the weakest performer. The order of vectors based on cell entry efficiency was overall similar in GFP^+^ human cells, with the exception that the relative contribution of AAV2 decreased substantially, suggesting that AAV2 transduced human cells but was less efficient at driving transgene expression. We also observed a relative drop for AAV3b and a corresponding increase in signal for AAV-NP59 (Fig 2c).

Looking at functional transduction (RNA/cDNA) of human hepatocytes in the FRG model, AAV-NP59 performed best in GFP^+^ human hepatocytes, with AAV-LK03, AAV3b/AAV8, AAV2 and AAV5 following behind.

The results for vector entry into macaque hepatocytes engrafted in FRG mice (RhFRG and CyFRG) were very similar to those obtained in humanized mice with AAV-NP59 being the serotype most efficient at cell entry (Fig. 2b). However, in bulk macaque hepatocytes, AAV-NP59 was followed by AAV8 (instead of AAV-LK03), AAV2 (instead of AAV3b), AAV-LK03, AAV3b and AAV5 (Fig. 2b). When analyzing GFP^+^ cells, we observed that highly performing variants AAV-NP59, AAV8 and AAV-LK03 showed an overall gain in contribution at the entry level, performance of AAV3b did not change substantially, while AAV2 and AAV5 showed a drop in efficiency compared to unsorted cells (Fig. 2c). Data from analyzed RNA confirmed this trend with AAV-NP59 showing by far the strongest contribution in simian cells sorted for GFP expression. AAV8 was the second-best performing variant and AAV-LK03, AAV3b, AAV2 and AAV5 variants had lower contributions at the transcriptional level (Fig. 2d).

Expectedly, this experiment indicated that the analysis of vector performance at the DNA level using cells sorted for transgene expression (GFP^+^) is more closely aligned with the analysis at the RNA level than when analyzing DNA from bulk human/NHP hepatocytes. However, bulk DNA data is useful to gain insight into the cell-vector interactions that are not leading to strong transgene expression. The analysis at the DNA level revealed that xenograft models show the same trend regarding AAV5’s performance in human and NHP hepatocytes. The *in vivo* xenograft data indicated that AAV5 interacts with, or enters, NHP hepatocytes relatively well but encounters some intracellular block, preventing it from efficiently completing all necessary steps that lead to transgene expression.

The high performance of AAV-NP59 in hFRGs was expected based on previously generated data that showed a strong advantage over the five other AAVs used in this study.^35^ The fact that we see a similar trend for the NHP-repopulated FRG mice was a very interesting finding which may indicate a high performance of this variant in human/primate hepatocytes or a particularly xenograft-specific high performance.

### Non-human primate *in vivo* transduction

In the next part of the study, we compared the performance of the six vectors *in vivo* in a widely accepted preclinical model of human liver, the NHP (Fig. 3a). To enable studies of multiple vectors in the same immunocompetent animal, the cynomolgus monkey underwent immunoadsorption (antibody depletion) to reduce antibody concentration, as previously described.^46^ Following this treatment, the NHP was infused with 4.2 × 10^13^ vg total (1.3 × 10^13^ vg/kg; 7.0 × 10^12^ vg/variant) of the barcoded AAV mix. The animal was sacrificed one week after systemic vector infusion and 21 different tissues were harvested and processed for downstream analysis.

**Figure 3.**
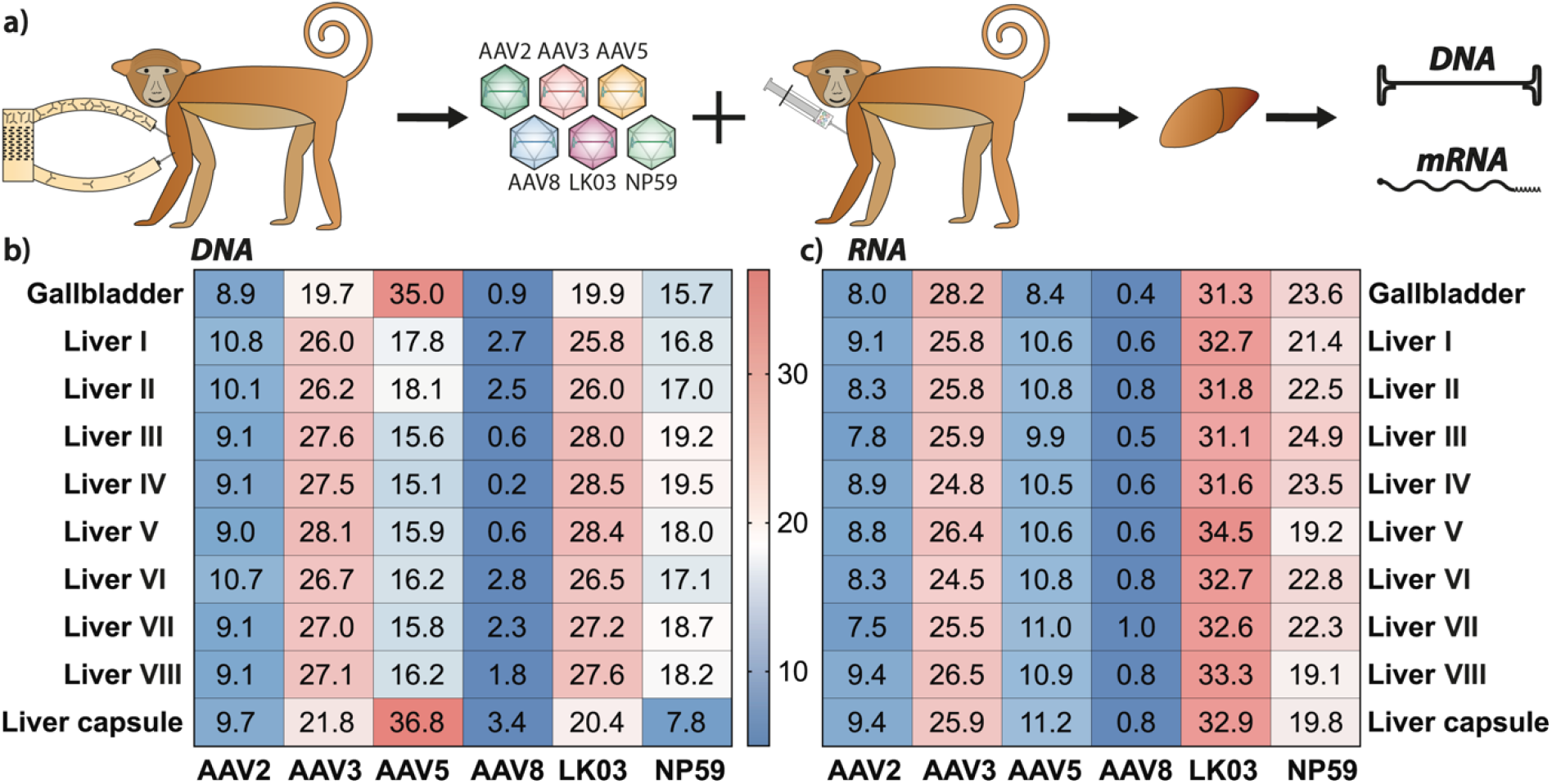
*In vivo* results cynomolgus monkey liver. (a) Schematic of column-based antibody depletion followed by non-human primate transduction. (**b**) NGS read contribution (%) for each AAV from whole tissue extracted DNA. (**c**) NGS read contribution (%) for each AAV from whole tissue mRNA-derived complementary DNA.

The samples were analyzed for vector copy number using ddPCR and transgene expression using reverse-transcriptase-ddPCR (RT-ddPCR) (Supplementary Table 2, Fig. 4b). Since the liver was the main organ of interest in this study, samples were taken from eight different regions of the liver (see Supplementary Fig 4a for indication of the liver regions analyzed) as well as from the gallbladder and the liver capsule and were analyzed for individual vector performance at the cell entry (DNA) and transgene expression (RNA/cDNA) levels using NGS. As expected based on previous publications,^35,54^ and the fact that all six vectors studied are known to be liver-tropic, vector copy number analysis showed that at the dose used the liver, gallbladder, and spleen were the organs with the highest levels of transduction (Supplementary Fig. 4b).

Non-liver organs appeared to be most efficiently entered by AAV5 (Supplementary Fig. 4c). Whether these results are truly reflecting the ubiquitous activity of AAV5 or are an artefact of the very low vector copy number in these organs cannot be inferred from this dataset. Unsurprisingly, given that the GFP transgene expression was driven by the ApoE/hAAT liver-restrictive promoter, transgene expression could only be detected in the liver and gallbladder (Supplementary Fig. 4b).^36,55,56^

NGS analysis of NHP liver samples showed that AAV-LK03 and AAV3b were the most effective variants at transducing most regions of the liver. AAV-NP59 was the next best performer, followed closely by AAV5. Interestingly, AAV5 was the serotype that entered cells most efficiently in the gallbladder and liver capsule (Fig. 3b). AAV2 and AAV8 were the least efficient serotypes in terms of cell entry (Fig. 3b). In terms of transgene expression, AAV-LK03 was the best performing variant, followed by AAV3b and AAV-NP59. AAV5, AAV2 and AAV8 performed relatively poorly (Fig. 3c).

From previously published data, we were not surprised by AAV-LK03’s high performance^54^ and it was interesting to see how much better AAV-NP59 performed compared to the prototypical AAV2, which is highly homologous at the protein level, but potentially tissue culture-adapted,^42^ indicating that the previously published advantage of lower HSPG binding is beneficial in *in vivo* xenograft mouse models as well as NHP models.^31^ What was surprising, however, was the very low AAV8 performance,^54^ which warranted further investigation.

### Non-human primate serum analysis

Driven by the fact that AAV8 performance in the NHP liver was lower than anticipated based on published data,^54^ we analyzed the levels of anti AAV NAbs in the serum harvested prior to AAV infusion. Additionally, the clearance of AAVs from the serum was quantified. Both seroreactivity and AAV clearance were evaluated at five time points (Fig. 4a).

**Figure 4.**
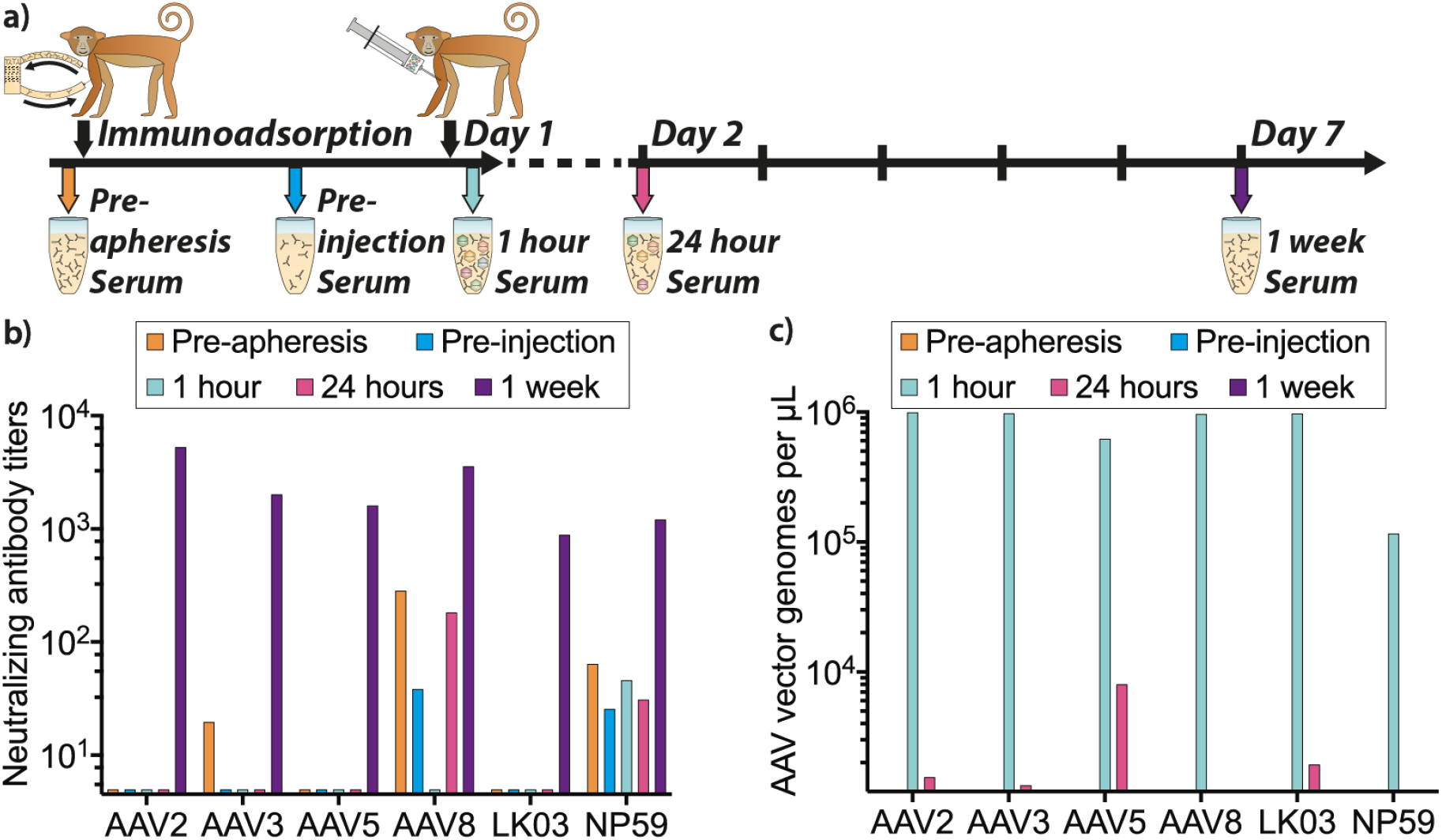
AAV-treated NHP serum analysis. (**a**) Schematic of serum collection before apheresis (antibody depletion), prior to AAV injection, one and 24 hours after injection as well as at sacrifice one week after injection. (**b**) Neutralizing antibody titers in collected serum at indicated time points for the indicated AAV variants. (**c**) AAV copy number per microliter and AAV variant from collected serum at indicated time points.

The neutralizing antibody titers showed that the NHP had pre-existing neutralizing antibodies (NAb) against AAV3b, AAV8 and AAV-NP59 prior to apheresis. These NAb titers were reduced by the apheresis in all cases. While the anti-AAV3 NAbs were effectively removed, antibodies against AAV8 and AAV-NP59 were not fully eliminated (Fig. 4b).

Seven days after vector administration we detected high NAb titers against all AAV variants (Fig. 4b). Previous publications stated that even low NAb titers against AAV8 capsids could have a strong neutralization effect *in vivo* in NHPs, and that this effect was substantially greater than expectations based on results from *in vitro* neutralization assays.^57,58^ Thus the presence of residual anti-AAV8 NAbs can potentially explain the low performance of AAV8 vector *in vivo*.

Analysis of AAV vector genomes in serum confirmed that substantial levels of AAV vectors were in circulation one hour after the injection, and that most of the vectors were cleared within the first 24 hours post-infusion. Particle concentrations were very similar between all variants apart from AAV-NP59, which showed an almost 10-fold lower concentration in the serum at the 1-hour time point compared to all other variants (Fig. 4c), potentially indicating a faster uptake of AAV-NP59 by the liver, uptake by other tissues, or an uptake by certain immune cells due to the observed interaction with NHP serum (Fig. 4b).

### The effect of NHP serum on the transduction of AAV vectors in xenograft models of the human liver

Next, we wanted to take advantage of the humanized and “monkeynized” FRG models to investigate the potential impact of the anti-AAV neutralizing antibodies in the NHP serum on vector transduction of primary hepatocytes. Using methods previously described,^39^ the equimolar mix of AAVs was co-incubated at a range of dilutions with NHP serum collected after the apheresis but before vector administration (Fig. 4a, “pre-injection serum”), and thus contained small titer of anti-AAV8 and anti-AAV-NP59 NAbs (Fig. 4b). The serum-AAV mix was subsequently injected into FRG mice repopulated with primary hepatocytes from either Rhesus macaque, Cynomolgus macaque or human origin (Fig. 5a). Human and NHP hepatocytes, as well as murine hepatocytes, were recovered from livers harvested seven days after systemic administration of the serum-AAV mix and individual vector transduction was analyzed at the DNA level using NGS of the barcoded genomic region.

**Figure 5.**
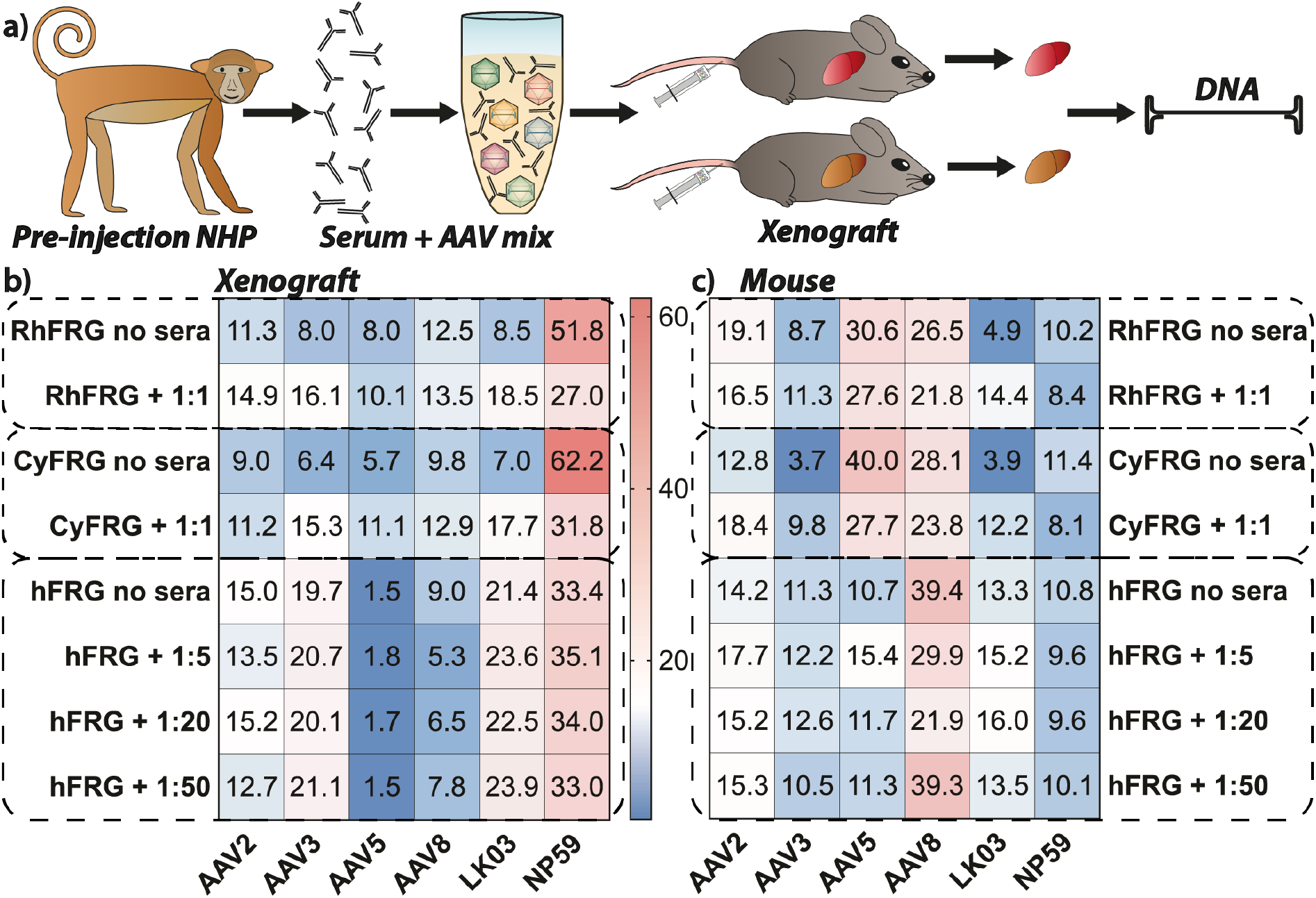
*In vivo* transduction with NHP pre-injection serum-incubated AAV mix. (**a**) Schematic of transduction of engrafted FRG mice with co-incubated AAV mix and antibodies from the post-apheresis/pre-injection step of the experiment using *Macaca fascicularis* NHP. (**b**) NGS read contribution (%) for each AAV from extracted xenograft DNA in absence or presence of serum in indicated dilutions. Cells were sorted for xenograft species. (**c**) NGS read contribution (%) for each AAV from mouse extracted DNA in absence or presence of serum in indicated dilutions. Cells were sorted for mouse origin. All data shown as ‘Rh/Cy/hFRG no sera’ are also used for Figure 2b and are shown again for ease of comparability.

AAV-NP59 was the top performer at the DNA level in FRG mice repopulated with Rhesus and Cynomolgus hepatocytes with and without vector pre-incubation with serum. However, the AAV-NP59 contribution in the group treated with the serum was substantially lower than in the group transduced with untreated vectors (Fig. 5b) indicating that the anti-AAV-NP59 antibodies were partially neutralizing the vector and thus decreasing its performance. Interestingly, there was no detected drop in the transduction efficiency of AAV8 following incubation with NHP sera. However, it is important to note that NGS percentages are relative to one another and thus the substantial drop in contribution from AAV-NP59 could mask a potentially reduced efficiency of neutralized AAV8. Indeed, relative transduction of all other vectors increased as the contribution of AAV-NP59 decreased, with AAV8 and AAV2 showing the lowest increase in transduction. This could indicate that the sera contained neutralizing antibodies against those two variants.

Interestingly, studies in FRG repopulated with human hepatocytes (hFRG) showed that performance of AAV-NP59 was not affected by the anti-AAV NAbs-containing serum and neither was the performance of AAV2, AAV3b, AAV5 or AAV-LK03, but the performance of AAV8 was reduced at the higher serum concentrations (Fig. 5b). All data shown as ‘Rh/Cy/hFRG no sera’ are also used in Figure 2b and are shown again for ease of comparability.

Lastly, analysis of the mouse hepatocytes recovered from the chimeric livers showed a mild drop in the performance of AAV8 and AAV-NP59 for NHP and human-repopulated mice (Fig. 5c). Interestingly, performance of AAV3b and AAV-LK03 in “monkeynized” FRGs appeared to have improved slightly following pre-incubation with NHP sera. While this can be partially explained by the previously mentioned fact that NGS reads for each vector are relative to one another, the effect could also indicate an active interaction between AAV3-like capsids with components of the NHP serum, as similar findings for some AAVs have been reported for interactions with human sera in mice.^59^ It is also important to note that the immunodeficient FRG mouse model may not fully recapitulate what happens to AAV-antibody-complexes in immunocompetent NHPs *in vivo,* as many immune cells are not fully developed.

### Seroprevalence of neutralizing antibodies against capsids used in this study

As transduction efficiency of an AAV capsid can be negatively affected by neutralizing antibodies, pre-existing immunity can exclude patients from an AAV gene therapy trial or clinical treatment.^60^ Therefore, we assessed the seroprevalence of NAbs against AAV variants used in this study in 85 human samples. Samples were collected from individuals younger than one year old (17 %), aged one to five years (35 %), six to ten years (16 %), eleven to twenty years (12 %), and older than twenty years old (20 %). The overall seroprevalence of NAb ranged from 14% (AAV5) to 29% (AAV3b) (Fig. 6a).

**Figure 6.**
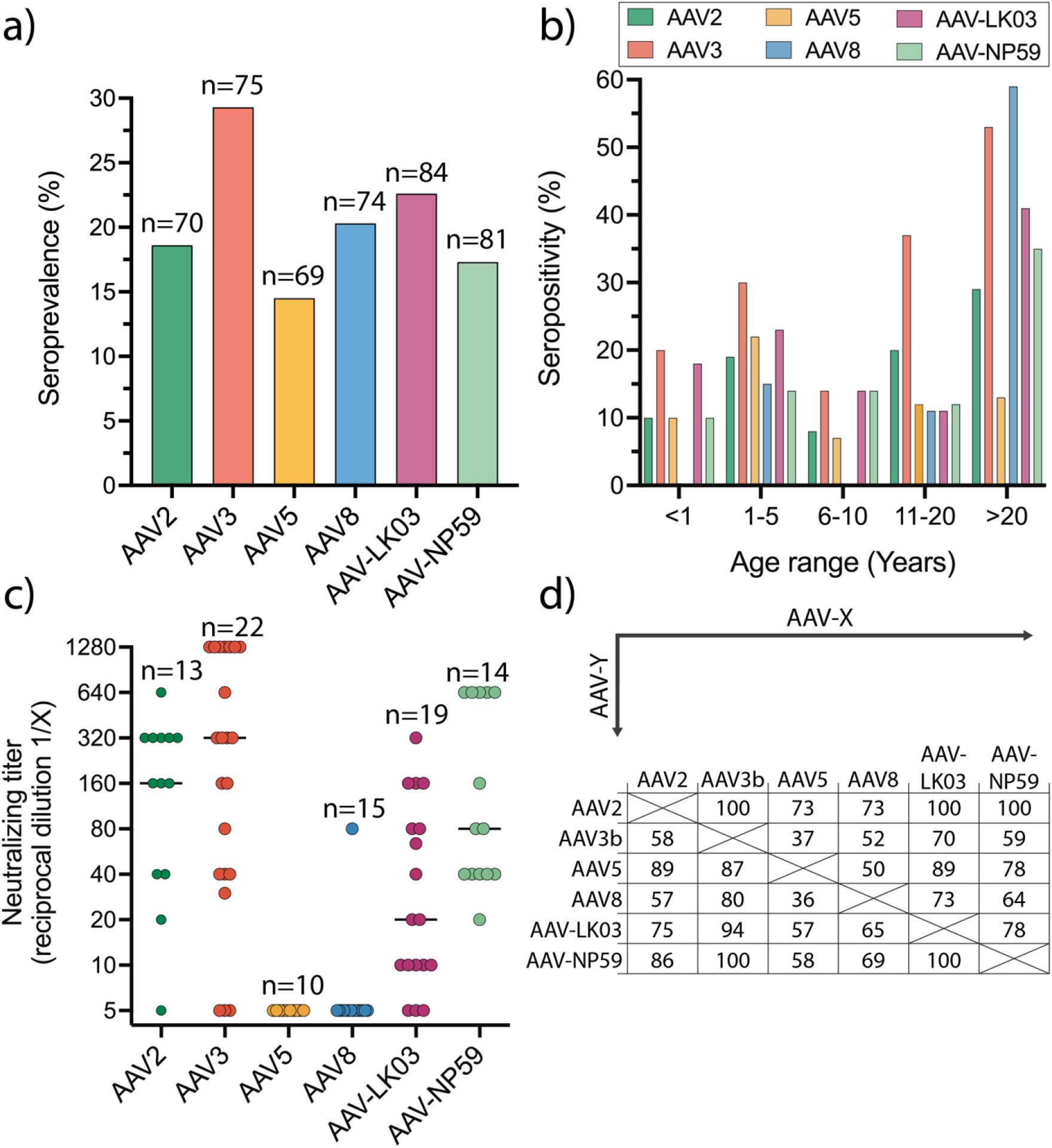
Seroprevalence and titers of neutralizing antibodies of liver-tropic capsids. (**a**) Seroprevalence per AAV serotype. (**b**) Seroprevalence according to age. (**c**) Neutralizing titer per serotype. Each dot represents a seropositive sample. The line represents the median. (**d**) Cross-reactivity by AAV serotype. Shown are the percentage of samples that are positive for AAV-X in samples that were positive for AAV-Y.

Seroprevalence was low during the first years of age and increased after the age of 10 years (Fig. 6b). NAb titers are higher for AAV3b, AAV2 and AAV-NP59 with median of 1/320, 1/160 and 1/80, respectively. Lower NAb titers were observed for AAV5, AAV8 and AAV-LK03 with medians of 1/5, 1/5 and 1/20, respectively (Fig. 6c). Strong cross-reactivity with other liver-tropic capsids tested was detected for AAV2, AAV-NP59 and AAV5 (Fig. 6d).

In the cohort of plasma samples tested for seroprevalence, 80% were from persons aged 20 years old or younger. This work corroborates previous findings that most pediatric individuals have not yet developed anti-AAV antibodies, which emphasizes a decisive immunological advantage of targeting this age group.^61,62^ The seroprevalence rates increased rapidly in teenage and adulthood where NAb seroprevalence rates could be as high as 80% for the some AAV serotypes.^27,62–66^ Our data for sera from the adult population showed lower neutralization rates compared to other seroprevalence studies^64^ but the findings were in accordance with our previous work in the British population.^45^ AAV2, AAV3b and surprisingly AAV-NP59 showed higher titers compared with AAV5, AAV8 and AAV-LK03, although the use of a higher dose for AAV5 and AAV8 may have reduced the sensitivity and partly underestimated the titres.^67^

Neutralizing antibody cross-reactivity was high between some liver-tropic capsids.^45,62,64^ These findings support the need for innovative immunosuppression protocols or antibody reduction methods to allow successful transduction in preimmunized patients and reinjections if needed.^60,68–70^

### Conclusions

Expectedly, we observed that the interaction of the AAV vectors and the target hepatocytes differed in *in vitro* and *ex vivo* cells as well as xenograft models and *in vivo* NHP transductions. However, even though a perfectly predictive preclinical model does not exist, our study shows that each model provides a unique insight into the vector function. However, without the benefit of clinical data, where each of the vectors would be tested for the delivery of the same transgene cassette to the liver, it is impossible to determine, with a high level of certainty, which of the models is the most predictive of human outcome.

Based on our results and the fact that each model brings a unique perspective that adds to the overall functional evaluation of AAV vectors, we propose that multiple models should be used to paint a more complete picture and help us make the most informed decision as to which vector should be used in each clinical application. To do so, it is critical to understand the strengths and weaknesses of each model. Specifically, our data confirmed that many tissue culture models are overly dependent on strong AAV binding to HSPG, which may not be directly applicable to an *in vivo* setting.^31,42^

We also found that AAV5 generally performed better in primary NHP hepatocytes *ex* and *in vivo* compared to human hepatocytes in the same experimental settings. As this finding was consistent between models, it may explain the lower-than-expected outcomes in several clinical trials using AAV5 to target the human liver.^24,25^ As AAV8 performed better in NHP-FRGs than in hFRGs, our data could suggest that a similar mechanism could affect AAV8’s performance in human hepatocytes.

Generally, our study showed that the FRG mouse model offers high flexibility and utility as it can be repopulated with primary hepatocytes from human and NHPs.^71^ Thus, this xenograft model allows investigators to gain a unique insight into the cross-species transferability of the AAV performances data from NHPs, the most sought after pre-clinical model of human liver, to human patients. Furthermore, our data showed that the correlation between data obtained from NHP and xenograft models was not perfect. This could be due to complex, and not fully understood, interactions between xenograft and mouse hepatocytes, competition in AAV uptake between the species, or the absence of a complete immune system in the mice.

However, the NHP model may not be perfect either. Apart from ethical considerations around studies involving NHPs, availability and cost can be prohibitive. The cost and availability of NHPs is also affected by the fact that animals need to be screened for pre-existing immunity as well as the fact that outbred NHPs are used, requiring many animals per group to account for natural differences between “subjects”. Based on the data presented, our current understanding of the individual preclinical models, as well as our insights into the AAV-cell interactions, we propose that initial studies of liver-targeting AAVs should include hFRG mice in the presence of human serum and/or pooled IgG before initiating studies involving large animals.

Based on data from a Hemophilia A clinical trial for AAV-LK03^30^ as well as data from this study (good efficiency at transducing NHP and human hepatocytes *ex vivo* and *in vivo* in the xenograft model and the NHP, and relatively low pre-existing immunity in the general population), we anticipate that the next generation bioengineered AAV-LK03 and AAV-NP59 vectors will be strong clinical candidates for liver targeted therapies.

## Acknowledgements

This work was supported by project grants from the Australian National Health and Medical Research Council (NHMRC) to L.L. and I.E.A. (APP1108311, APP1156431 and APP1161583) and Paediatrio Paediatric Precision Medicine Program to L.L. (PPM1 K5116/RD274). Work presented in Figures 3 and 4 were supported by funding from LogicBio Therapeutics. L.L. was also supported by research grants from the Department of Science and Higher Education of Ministry of National Defense, Republic of Poland, (“Kościuszko” k/10/8047/DNiSW/T – WIHE/3) and from the National Science Centre, Republic of Poland (OPUS 13) (UMO-2017/25/B/NZ1/02790). The work of I.E.A. was also supported by an Australian Research Council (ARC) Discovery Project (DP150101253). A.J.T. was supported by funding from The Wellcome Trust (grant no. 217112/Z/19/Z AJT).

This work was supported by funding to JB from the NIHR Great Ormond Street Hospital Biomedical Research Centre; Medical Research Council Grant/Award Number: MR/T008024/1; National Institute for Health Research; Innovate UK Biomedical Catalyst Early stage award 14720; Nutricia Metabolic Research Grant; London Advanced Therapy / Confidence in Collaboration award 2CiC017. The views expressed are those of the author(s) and not necessarily those of the NHS, the NIHR or the Department of Health.

## Author contributions

A.W., M.C-C., E.Z., D.S.G., D.P.P., S.A.W., J.B., G.G-A., and L.L. designed the experiments. A.W., M.C-C., K.L.D., E.Z., D.S.G., R.G.N., S.S., M.K., A.M., S.F., D.P.P., B.M.T., E.V., S.L.W., S.A.W., and L.L. generated reagents, protocols, performed experiments, and analyzed data. A.W., A.K.A., and L.L. wrote the manuscript and generated the figures. All authors reviewed, edited, and commented on the manuscript.

## Author Disclosure Statement

L.L., I.E.A. and A.J.T. have commercial affiliations. L.L. and I.A.E. have consulted on technologies discussed in this paper. L.L. and I.A.E. have stock and/or equity in companies with technologies broadly related to this study. L.L. is a co-inventors of, and receives licensing royalties from, several AAV variants used in the study. A.L.M. and T.E.H. are employees, shareholders, and/or optionees of Inventia Life Science Pty. Ltd. Inventia has an interest in commercializing the 3D bioprinting technology. All other authors declare no competing financial interests.

