## Supplemental Information for "Assessment of pre-clinical liver models based on their ability to predict the liver-tropism of AAV vectors"

**Supplementary Table 1. Components of human liver organoid expansion media**

| **Components** | **Concentration** | **Supplier** |
|---|---|---|
| Basal media:<br>Advanced DMEM/F-12 media<br>Penicillin-Streptomycin<br>HEPES<br>GlutaMAX | 86%<br>Neat<br>1x<br>10 mM<br>1x | <br>Life Technologies 12634-028<br>Life Technologies 15140-122<br>Life Technologies 15630-056<br>Life Technologies 35050-061 |
| Rspo1-conditioned media | 10% | In-house. See reference 71 |
| B-27 Supplement, without vitamin A | 1x | Life Technologies 12587-010 |
| N-2 Supplement | 1x | Life Technologies 17502-048 |
| Nicotinamide | 10 mM | Sigma N0636 |
| Human EGF | 50 ng/ml | Peprotech AF-100-15 |
| [$Leu^{15}$]-Gastrin I human | 10 nM | Sigma G9145 |
| N-acetylcysteine | 1 mM | Sigma A0737 |
| Human FGF-10 | 100 ng/ml | Peprotech 100-26 |
| Human HGF | 25 ng/ml | Peprotech 100-39 |
| Forskolin | 10 µM | Tocris Bioscience 1099 |
| A83-01 | 5 µM | Tocris Bioscience 2939 |

**Supplementary Table 2. Primers used in the presented study**

| **Name** | **Sequence (5’->3’)** | |
|---|---|---|
| **ddPCR** | **Sequence (5’->3’)** | |
| eGFP_F | TCAAGATCCGCCACAACATC | |
| eGFP_R | TTCTCGTTGGGGTCTTTGCT | |
| NHP_alb_F | CGCAACTCTTCGTGAAACCTATGG | |
| NHP_alb_R | CACATCAACCTCTGGTCTCACC | |
| NHP_b-actin_F (RT) | CAACGAGCGGTTCCGCTG | |
| NHP_b-actin_R (RT) | CAGCACTGTGTTGGCGTACAG | |
| **cDNA synthesis** | **Sequence (5’->3’)** | |
| WPRE_R | GGATTTATACAAGGAGGAGAAAATGAAAG | |
| **Next-generation sequencing** | **Primer barcode (5’->3’)** | **Main oligo sequence (5’->3’)** |
| GFP_BC_WPRE_F01 | GTTCA | GCTGGAGTTCGTGACCGCCG |
| GFP_BC_WPRE_F02 | GTCAT | GCTGGAGTTCGTGACCGCCG |
| GFP_BC_WPRE_F03 | CTGTA | GCTGGAGTTCGTGACCGCCG |
| GFP_BC_WPRE_F04 | GTATT | GCTGGAGTTCGTGACCGCCG |
| GFP_BC_WPRE_F05 | CTAGT | GCTGGAGTTCGTGACCGCCG |
| GFP_BC_WPRE_F06 | ACTTC | GCTGGAGTTCGTGACCGCCG |
| GFP_BC_WPRE_F07 | CCTAT | GCTGGAGTTCGTGACCGCCG |
| GFP_BC_WPRE_F08 | ACTGA | GCTGGAGTTCGTGACCGCCG |
| GFP_BC_WPRE_F09 | TCCAA | GCTGGAGTTCGTGACCGCCG |
| GFP_BC_WPRE_F10 | GCATT | GCTGGAGTTCGTGACCGCCG |
| GFP_BC_WPRE_F11 | TCAAG | GCTGGAGTTCGTGACCGCCG |
| GFP_BC_WPRE_F12 | TCAGA | GCTGGAGTTCGTGACCGCCG |
| GFP_BC_WPRE_F13 | TCGTA | GCTGGAGTTCGTGACCGCCG |
| GFP_BC_WPRE_F14 | ACGAT | GCTGGAGTTCGTGACCGCCG |
| GFP_BC_WPRE_F15 | TATCC | GCTGGAGTTCGTGACCGCCG |
| GFP_BC_WPRE_F16 | CATGA | GCTGGAGTTCGTGACCGCCG |
| GFP_BC_WPRE_F17 | CACTC | GCTGGAGTTCGTGACCGCCG |
| GFP_BC_WPRE_F18 | GACTA | GCTGGAGTTCGTGACCGCCG |
| GFP_BC_WPRE_F19 | TACCC | GCTGGAGTTCGTGACCGCCG |
| GFP_BC_WPRE_F20 | GACAT | GCTGGAGTTCGTGACCGCCG |
| GFP_BC_WPRE_F21 | GAAGA | GCTGGAGTTCGTGACCGCCG |
| GFP_BC_WPRE_R | CAACATAGTTAAGAATACCAGTCAATCTTTCAC | |

**Supplementary Table 3. Top 20 up- or down-regulated genes in 3D-cultured hepatocytes in printed hydrogels**

| gene | name | up/ down | gene type | function | antisense to | antisense function | TMM fold change |
|---|---|---|---|---|---|---|---|
| **ENSG00000253438.4** | **PCAT1** | **up** | **lncRNA** | **Inhibiting p53/ upregulating myc** | **N/A** | **N/A** | **53.37158** |
| ENSG00000224251.6 | AL391427.1 | up | lncRNA | unknown | AKR1C2 | Aldo-keto Reductase | 23.10789 |
| ENSG00000227227.1 | AC017101.1 | up | lncRNA | unknown | ITGAV | Integrin sub-unit | 22.62667 |
| ENSG00000243415.2 | AC107021.1 | up | lncRNA | unknown | PLOD2 | Collagen synthesis | 16.44496 |
| ENSG00000120725.13 | SIL1 | up | protein_coding | Nucleotide exchange factor | N/A | N/A | 14.0066 |
| ENSG00000227110.7 | LMCD1-AS1 | up | lncRNA | Antisense RNA | LMCD1 | Zinc-finger, transcription cofactor | 13.30003 |
| ENSG00000145439.12 | CBR4 | up | protein_coding | Fatty acid biosynthesis | N/A | N/A | 13.15157 |
| **ENSG00000183098.11** | **GPC6** | **up** | **protein_coding** | **Heparan sulfate proteoglycan** | **N/A** | **N/A** | **12.47776** |
| ENSG00000225205.5 | AC078883.1 | up | lncRNA | unknown | ITGA6 | Integrin sub-unit | 12.03583 |
| ENSG00000254733.1 | AP001831.1 | up | lncRNA | unknown | ME3 | Malate dehydrogenase | 11.27253 |
| ENSG00000278518.1 | AL161645.1 | down | lncRNA | unknown | CYP2E1 | Monooxygenase | 0.01906202 |
| ENSG00000246090.7 | AP002026.1 | down | lncRNA | unknown | ADH | Alcohol dehydrogenase | 0.01927675 |
| ENSG00000237037.9 | NDUFA6-DT | down | lncRNA | unknown | NDUFA6 | Mitochondrial membrane respiratory chain | 0.01997939 |
| ENSG00000146215.13 | CRIP3 | down | protein_coding | inflammation/cancer | N/A | N/A | 0.0250615 |
| ENSG00000012061.15 | ERCC1 | down | protein_coding | Nucleotide excision repair | N/A | N/A | 0.02817534 |
| ENSG00000251139.2 | AC084871.1 | down | lncRNA | unknown | ACSL1 | Fatty acid biosynthesis | 0.02820471 |
| ENSG00000268895.6 | A1BG-AS1 | down | lncRNA | Antisense RNA | A1BG | Plasma glycoprotein | 0.03172232 |
| ENSG00000198739.11 | LRRTM3 | down | protein_coding | Synaptic acivity | N/A | N/A | 0.03399672 |
| ENSG00000231424.3 | BX284613.2 | down | lncRNA | unknown | FMO1 | Monooxygenase | 0.03403329 |
| ENSG00000255240.6 | AP001636.3 | down | lncRNA | unknown | GLYATL1 | Acyltransferase | 0.03589633 |

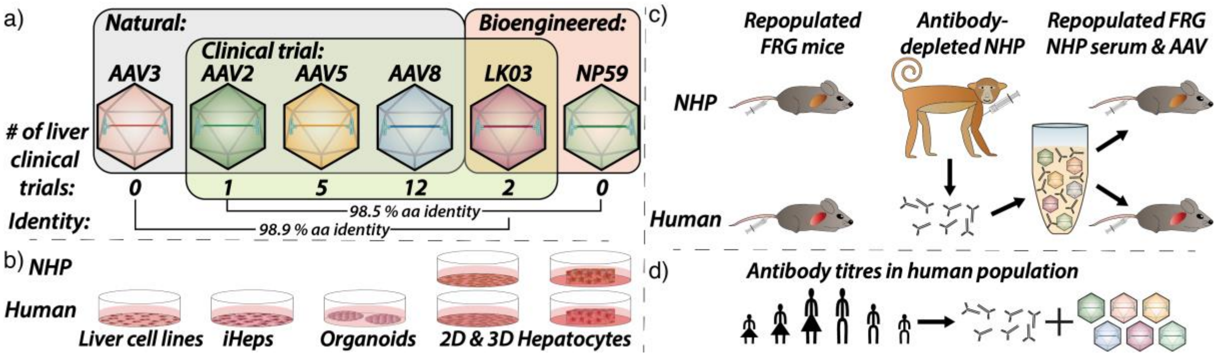


**Supplementary Figure 1. Overview of the hepato-tropic AAV and pre-clinical liver model comparison.** (**a**) AAVs chosen for this study based on their use in clinical trials or expected high performance in human hepatocytes based on previous studies and their amino acid identity. (**b**) Human and simian *in vitro* and *ex vivo* models of hepatocytes. (**c**) Human and simian *in vivo* models of hepatocytes. (**d**) Evaluation of antibody titres in human population for all six AAV variants included in the study. Abbreviations: AAV: adeno-associated virus; aa: amino acid; FRG: *Fah-/-/Rag2-/-/Il2rg-/-*; NHP: non-human primate

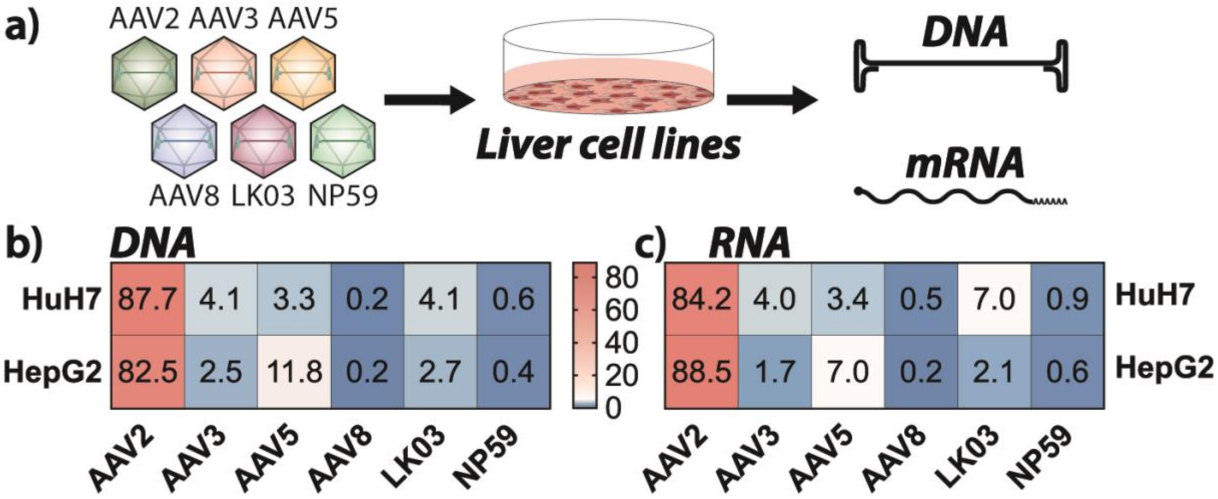


**Supplementary Figure 2. AAV performance in hepatocellular carcinoma cell lines.** (**a**) Schematic of transduction of indicated hepatocellular carcinoma cell lines. (**b**) NGS read contribution (%) for each AAV from extracted DNA. (**c**) NGS read contribution (%) for each AAV from mRNA-derived complementary DNA.

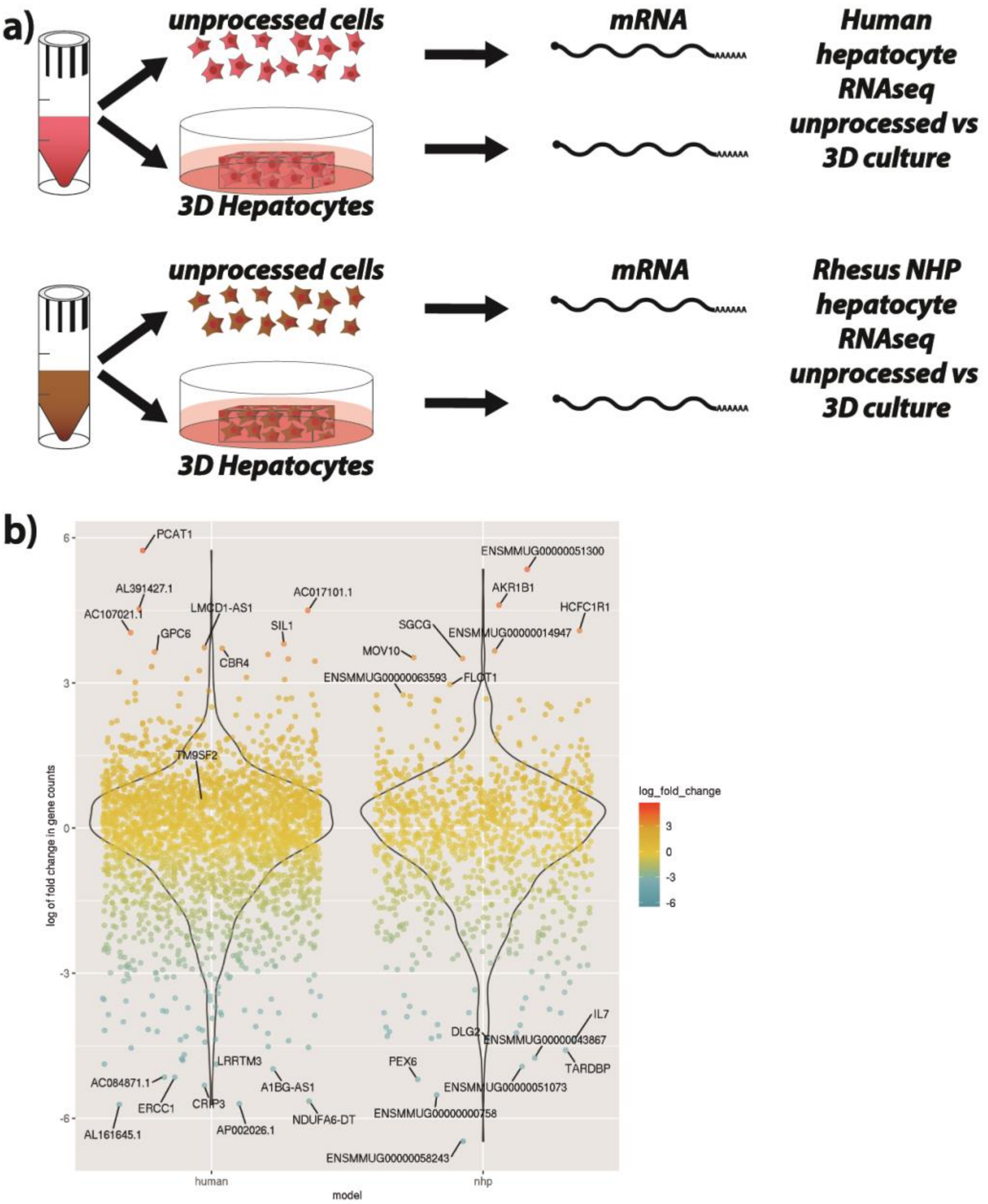


**Supplementary Figure 3. Transcriptomic differences in 2D and 3D-cultured hepatocytes.** (**a**) Schematic representation of experimental workflow. (**b**) Change in gene expression between normal and 3D cultured hepatocytes displayed as log of fold change.

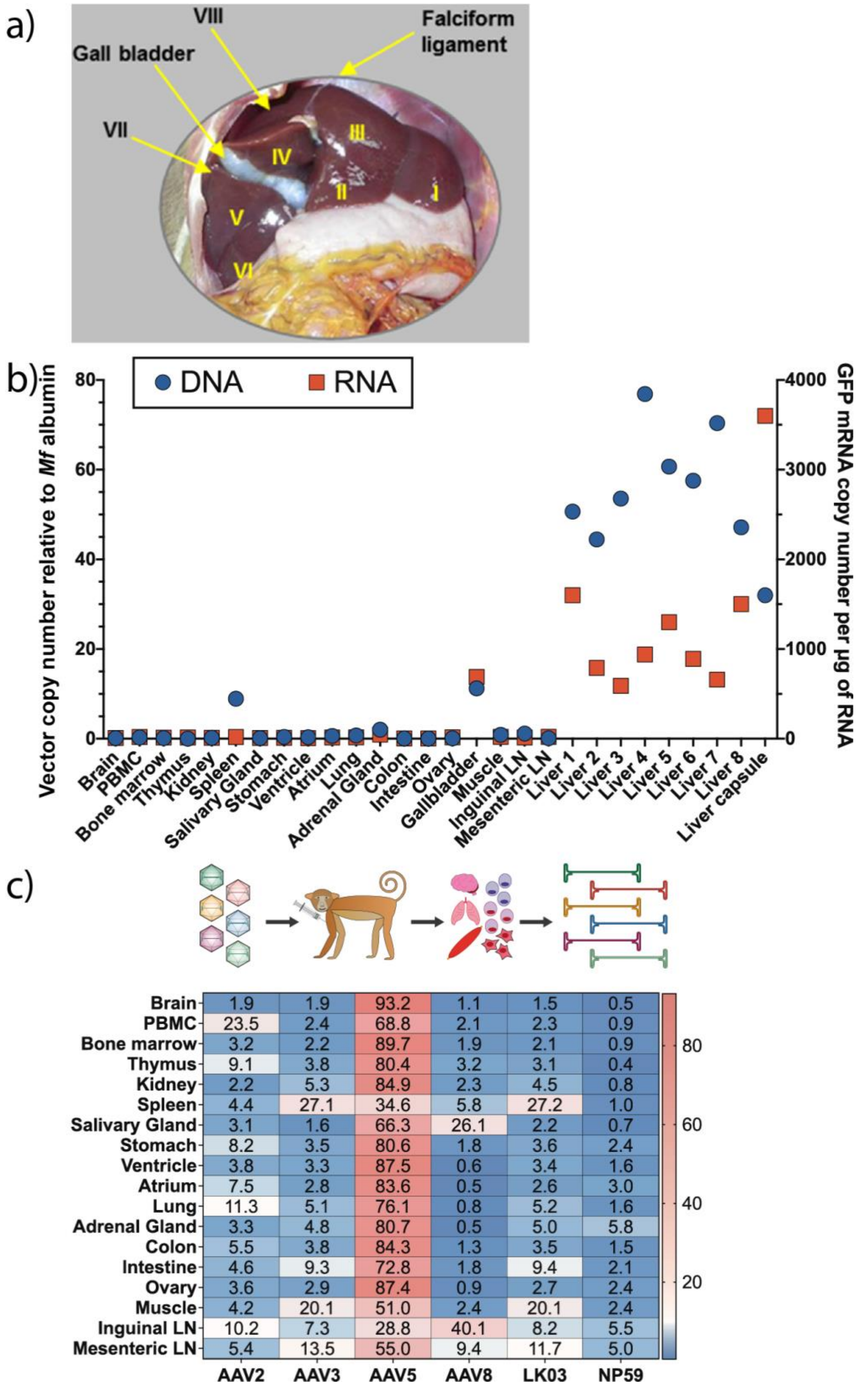


**Supplementary Figure 4. Cynomolgus monkey AAV biodistribution.** (**a**) Representation of liver regions indicated in **Figure 3**. (**b**) DNA (left y-axis) and mRNA/cDNA (right y-axis) vector copy number in the indicated organs and liver regions. (**c**) NGS read contribution (%) for each AAV from extracted DNA in the indicated cells and organs. Abbreviations: PBMC: peripheral blood-derived mono-nuclear cells; LN: lymph node
